# Chemical modulation of the unfolded protein response reveals an antiviral role for the PERK pathway in human coronavirus 229E infection

**DOI:** 10.1101/2025.09.11.675615

**Authors:** Isabella E. Pellizzari-Delano, Trinity H. Tooley, Keya Jani, Carla E. Gallardo-Flores, Che C. Colpitts

**Affiliations:** Department of Biomedical and Molecular Sciences, Queen’s University, Kingston, ON K7L 3N6

## Abstract

Broad spectrum antivirals are critical to respond rapidly to the threat posed by newly emerging RNA viruses. One potential candidate is the natural compound thapsigargin (Tg). Tg potently induces endoplasmic reticulum (ER) stress and activates the unfolded protein response (UPR). Recent studies have demonstrated that Tg has robust antiviral activity against several human coronaviruses (CoVs), including SARS-CoV-2, although the specific antiviral mechanism(s) have remained unclear. Here, we aimed to characterize the role of the UPR in the antiviral activity of Tg against HCoV-229E, a model common cold CoV. Consistent with previous findings, we show that a short 30-minute priming of A549 cells with Tg potently inhibits HCoV-229E infection. Time-of-addition assays showed that Tg is most effective when added up to 8 hours post-infection. Furthermore, Tg inhibits the accumulation of double-stranded RNA in infected cells, suggesting that Tg inhibits early stages of viral RNA replication. Using selective UPR pathway inhibitors to narrow down the role of these pathways in mediating the antiviral effect of Tg, we show that the inhibition of IRE1 or ATF6 does not impair the ability of Tg to inhibit HCoV-229E infection. The use of stable knockdown A549 cells in which IRE1, PERK, or ATF6 expression was silenced further revealed that the antiviral activity of Tg is not dependent on the expression of any of the three UPR sensors individually. However, HCoV-229E replication is inhibited in A549-shIRE1 cells, or in cells treated with the IRE1 inhibitor (KIRA6), suggesting that IRE1 activation may play a pro-viral role during HCoV-229E infection. Selective UPR pathway activators were used to further probe down the role of each pathway during HCoV-229E infection. Selective activation of the PERK pathway, but not IRE1 or ATF6 pathways, inhibits HCoV-229E infection. Lastly, to more broadly test the antiviral role of PERK against CoV RNA replication, we used BHK-21 cells that stably express a SARS-CoV-2 replicon. We show that selective PERK activation robustly inhibits SARS-CoV-2 replication, comparable to Tg. Overall, these findings provide insight into the antiviral mechanism(s) of Tg against CoV infection and demonstrate that modulation of the UPR may be exploited as an antiviral strategy.

## Introduction

Five years after the beginning of the COVID-19 pandemic, the significant global health threat posed by newly emerging viruses is still concerning. Although the COVID-19 pandemic demonstrated that effective vaccines and antiviral drugs can be developed for a newly emerging virus, there is a need for broadly acting, host-centric antiviral drugs that could be readily deployed in the context of a future, novel emergence event. Antivirals that do not target specific viral proteins, but rather function by enhancing host antiviral responses, are advantageous as they are expected to have broad antiviral activity and a high barrier to resistance^1^. One such compound that exhibits broad spectrum antiviral activity is thapsigargin (Tg).

Recent literature has demonstrated the broad-spectrum antiviral activity of Tg, a natural product isolated from the Mediterranean plant, *Thapsia garganica*. Indeed, Tg possesses robust and broadly acting antiviral activity against human coronaviruses (CoVs), including endemic viruses such as HCoV-229E and HCoV-OC43, as well as the highly pathogenic MERS-CoV and SARS-CoV-2^2,3^. Tg potently inhibits the replication of HCoV-229E, as marked by a reduction in the levels of viral non-structural proteins (nsp) 8 and 12, key components of the viral replication complex^3^. Interestingly, while both Tg and HCoV-229E infection induce ER stress in Huh7 cells, Tg-induced ER stress was proposed to reverse CoV-mediated inhibition of stress response factors, thereby protecting against infection^3^. Furthermore, Tg has been shown to inhibit the replication of other major respiratory viruses, including respiratory syncytial virus (RSV) and influenza A virus (IAV)^4^, in which the antiviral effect of Tg against IAV was accompanied by enhanced host antiviral interferon responses.

Tg is a well-known inhibitor of the sarcoplasmic/endoplasmic reticulum (ER) Ca^2+^ ATPase (SERCA) pump^5^, which plays a critical role in regulating Ca^2+^ homeostasis across the cell cytoplasm and ER lumen. Inhibition of SERCA by Tg depletes Ca^2+^ stores within the ER, inducing ER stress and the unfolded protein response (UPR)^5^. The UPR is a highly conserved mammalian response pathway that transcriptionally upregulates genes that function to restore proteostasis, or, under periods of intense and prolonged stress, induce cellular apoptosis^6^. The UPR is comprised of three signalling pathways, each regulated upstream by the activation of a corresponding transmembrane ER resident sensor/receptor, namely inositol requiring protein 1 (IRE1), PKR-like endoplasmic reticulum kinase (PERK), and activating transcription factor 6 (ATF6)^6^. The endonuclease activity of IRE1 mediates the unconventional splicing of target gene X binding protein 1 (XBP1). This splicing event generates a frameshift variant known as XBP1 spliced (XBP1s) that functions as a potent transcription factor that upregulates genes that promote protein folding and ER-associated degradation^7^. Activation of PERK leads to the phosphorylation of eukaryotic translation initiation factor 2α (eIF2α), which temporarily halts global cap-dependent translation, limiting the synthesis of new proteins entering the ER^8^. Phosphorylation of eIF2α mediates the activation of activating transcription factor 4 (ATF4), which controls the expression of genes involved in alleviating ER stress, or inducing apoptosis, such as including CCAAT-enhancer-binding protein homologous protein (CHOP)^9,10^. Lastly, following activation, ATF6 translocates to the Golgi, where it is proteolytically cleaved to reveal its NH_2_ domain, also known as ATF6-N^11^. ATF6-N also functions as a transcription factor that upregulates the expression of genes involved in restoring proteostasis^11^.

One hypothesis is that UPR activation mediates the antiviral effect of Tg. However, the exact antiviral mechanism(s) of Tg against CoVs remains unclear, and the role of specific UPR pathways in mediating the antiviral activity of Tg is not well understood. In this work, we sought to further our understanding of the antiviral mechanism of Tg against the model common cold CoV HCoV-229E by characterizing which branches of the UPR modulate HCoV-229E infection and could mediate the antiviral effect of Tg. We have supported previous findings that Tg potently inhibits HCoV-229E infection^3^. We have shown that Tg is most effective when added up to 8 hours post infection, suggesting that Tg treatment inhibits an early post-entry step of the viral replication cycle, such as viral protein expression or RNA synthesis. Activation of all three UPR pathways is expedited in cells primed with Tg. Additionally, chemical modulation of the UPR during infection using selective small molecule activators and inhibitors suggests that activation of the PERK pathway is antiviral against both HCoV-229E and SARS-CoV-2 replication. On the other hand, inhibition of IRE1 strongly inhibited HCoV-229E infection, suggesting a proviral role for the IRE1 pathway. We show that genetic silencing of IRE1, ATF6, or PERK expression does not impair the antiviral activity of Tg, suggesting that none of the UPR pathways individually mediate the antiviral effect of Tg. Overall, these findings demonstrate that pharmacological modulation of the UPR during infection may provide a host-centric antiviral strategy and provide insight into the antiviral mechanism of Tg against human CoV infection.

## Methods

### Cells and viruses

Parental A549 cells (BEI Resources NR-52268) were cultured in Ham’s F-12 K (Kaighn’s) medium (Fisher Scientific #21127030) supplemented with 10% fetal bovine serum (FBS) and 1% penicillin/streptomycin (pen/strep). Huh7 cells (JCRB0403) were obtained from the Japanese Collection of Research Bioresources Cell Bank. 293T/17 (CRL-11268) cells were obtained from American Type Culture Collection. Huh7 and 293T/17 cells were cultured in Dulbecco’s Modified Eagle Medium (DMEM; ThermoFisher #11995065) supplemented with 10% FBS and 1% pen/strep. BHK-21 SARS-CoV-2-Rep-NanoLuc-Neo cells^12^ were obtained from BEI Resources (NR-58876), and cultured in DMEM supplemented with 10% FBS, 1% pen/strep, and 200 μg/mL of G418 (ThermoFisher #10131027). HCoV-229E was obtained from BEI Resources (NR-52726) and propagated on Huh7 cells as previously described^13^.

### UPR inhibitors and activators

Thapsigargin (Tg) was kindly provided by Dr. Andrew Evans (Queen’s University) or purchased from Thermo Scientific (T7458). AA147 (E0745), IXA4 (S9797), CeapinA7 (E1099) and KIRA6 (S8658) were purchased from Selleckchem. CCT020312 (324879) and GSK2606414 (516535) were purchased from Sigma-Aldrich. Stocks were prepared in dimethyl sulfoxide (DMSO), which was used as the vehicle control.

### Plasmids

Lentiviral pLKO.1-shATF6, pLKO.1-shIRE1 and pLKO.1-shPERK constructs were kindly provided by Dr. Craig McCormick (Dalhousie University, Canada). For production of lentivirus, psPAX2 (#12260) and VSV-G (# 8454) were obtained from Addgene.

### Generation of stable cell lines

To produce lentivirus, 293T/17 cells were seeded in a 6-well plate at 8 x 10^6^ cells/well overnight. The next day, transfection mixes were prepared as followed: 600 ng psPAX2 packaging plasmid, 800 ng pLKO.1 transfer plasmid and 600 ng VSV-G envelope plasmid and 6 μL of lipofectamine 2000. Supernatants were harvested 48- and 72-hours post transfection and lentiviruses were filtered using a 0.45 µm filter. For the generation of shCTRL, shATF6, shIRE1 and shPERK cell lines, A549 cells were seeded in a 6-well plate at a density of 2 x 10^5^ cells/well overnight and subsequently transduced with lentivirus in the presence of 8 µg/mL of polybrene. Transduced cells were selected at 72 hours post transduction in either 3 µg/mL puromycin or 10 µg/mL blasticidin. Successful knockdown was confirmed by western blot analysis.

### Antibodies

Primary antibodies against IRE1α (Cell Signaling Technology (CST) 3294, dilution 1:1000), ATF6 (CST 8089, dilution 1:1000), PERK (CST 5683, dilution 1:1000), XBP1s (CST 12782T, dilution 1:1000), HERPUD1 (Abcam ab150424, dilution 1:1000), eIF2α-P (Ser-51) (CST 3398S, dilution 1:1000), HCoV-229E N (SinoBiological 40640-T62, dilution 1:10,000), beta-actin (Abcam ab8226, dilution 1:5000), alpha-tubulin (Thermo Scientific MA5-31466, dilution 1:5000), and GAPDH (Thermo Scientific MA5-15738, dilution 1:1000) were used for western blotting. The secondary antibodies Licor IRDye-680 (red) goat anti-mouse and IRDye-800 (green) goat anti-rabbit were also used. For IF, cells were stained using antibodies against calnexin (Sigma C4731, dilution 1:200) or dsRNA (Kerafast ES2001, dilution 1:65), followed by AlexaFluor-488 (CST 4408S or 4412S) or AlexaFluor-555 (CST 8890S or 8889S) conjugated secondary antibodies diluted 1:1000.

### Western blot

Protein was extracted from cell lysates using Laemmli buffer (12.5 mM Tris-HCl (pH 6.8), 4% SDS, 20% glycerol) supplemented with ETDA, protease and phosphatase inhibitors. Cell lysates were sheared using QIAshredder columns (QIAGEN, 79654). Samples were separated by SDS-PAGE and transferred to nitrocellulose membranes using the TransBlot Turbo System (Bio-Rad) as per the manufacturer protocol. Membranes were blocked in 5% milk dissolved in 1X TBS-T (Tris-buffered saline with 0.1% Tween-20) for 1 hour. Primary antibodies were diluted in blocking buffer and were incubated at 4°C overnight. The next morning, membranes were washed with 1X TBS-T and then incubated at room temperature for 1 hour with secondary antibodies diluted 1:10,000 in blocking buffer. Membranes were then washed with 1X TBS-T followed by 1X TBS and imaged using a Licor Odyssey® Clx imager.

### RT-qPCR

Cellular RNA was isolated from cellular lysates using the Monarch Total RNA Miniprep Kit (New England Biolabs) as per the manufacturer’s protocol. Extracted RNA was reverse transcribed to generate cDNA using the High-Capacity cDNA Synthesis Kit (Thermo Scientific). Quantitative PCR (qPCR) was performed using the PowerTrack^TM^ SYBR Green Master Mix (Thermo Scientific) with specific primers outlined in table 1. Viral and cellular gene expression was normalized to cellular actin using the 2^−dCt^ method.

**TABLE 1.**
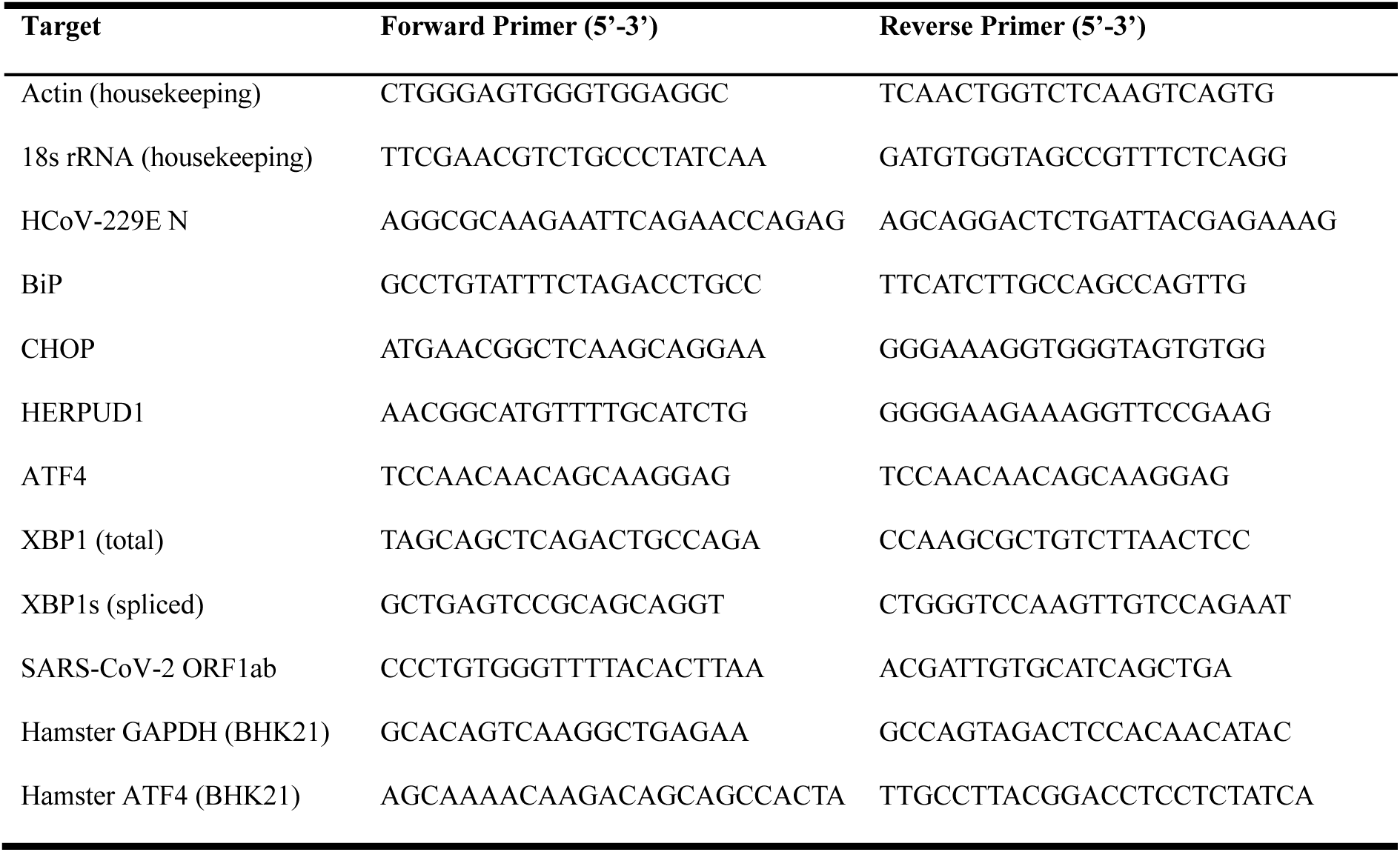

### Luciferase assays

Luciferase reporter activity in the BHK-CoV-2 replicon cells was measured using the NanoFuel® GLOW Assay for Oplophorus Luciferases from NanoLight Technology (Cat# 325). The NanoFuel® GLOW Reagent was brought to room temperature and 20 µL of 50x Oplo-GLOW Substrate was added to 1 mL of NanoFuel® GLOW Reagent. Once mixed, equal volume of the reagent-substrate mixture was added (100 µL in a 96-well plate) to each well containing 100 µL of media. The plate was incubated for 5 minutes after which the cells were scraped and transferred to a white flat-bottom 96-well plate for measurement using a Promega GloMax. For measurement of HCoV-229E Renilla luciferase reporter activity, 50 μL of prepared 1 mg/mL coelenterazine substrate (NanoLight) was injected into white plates containing 20 μL of cell lysate and measured by Promega GloMax.

### Immunofluorescence

A549 or BHK-CoV-2 replicon cells were seeded on coverslips in a 12-well plate at a density of 2 x 10^5^ cells/well and incubated overnight. A549 cells were subsequently infected with HCoV-229E at a multiplicity of infection (MOI) of 0.5 plaque-forming units (PFU)/cell for 24 hours. Cells were fixed in 10% formalin for 1 hour at room temperature. Fixed samples were permeabilized with 0.1% Triton-X 100 and blocked in PBS containing 5% FBS solution for 1 hour. Cells were stained for calnexin or dsRNA, followed by staining with AlexaFluor-488 or AlexaFluor-555 conjugated secondary antibody. Glass coverslips were mounted on microscope slides using FluoroMount G with DAPI (Thermo Scientific) and visualized at 40x magnification using a Nikon Eclipse Ts2-FL inverted microscope or 63x magnification under water immersion using a Leica Mica confocal microscope.

### Cell viability

The viability of A549 cells in the presence of UPR chemical modulators was assessed using the AlamarBlue reagent (Thermo Scientific) according to the manufacturer protocol.

### Plaque assay

Huh7 cells were seeded in a 12-well plate at 3.5 x 10^5^ cells/well in DMEM containing 10% FBS. Cellular supernatants containing HCoV-229E were serially diluted in serum-free DMEM. Cells were infected with diluted virus at 37°C for 2 hours. Viral inocula were removed and replaced with plaquing media (DMEM containing 1.2% carboxymethylcellulose and 2% FBS), and cells were incubated at 33°C for four days. Cells were fixed and stained using a 0.5% crystal violet solution in 20% methanol to visualize plaques.

### Statistical analysis

Statistical analyses were conducted using GraphPad Prism 10 version 10.5.0.

## Results

### Thapsigargin inhibits early stages of HCoV-229E RNA replication

Recently, Tg **(Figure 1A)** has been shown to possess potent and broadly acting antiviral activity against several unrelated viruses, including human coronaviruses^2^. While the antiviral mechanism of Tg is still unclear, its ability to induce ER stress and activate the UPR is well documented. Therefore, we hypothesized that activation of the UPR by Tg during viral infection mediates its antiviral activity. To confirm the inhibitory effect of Tg against HCoV-229E in our cell model, A549 lung epithelial cells were pre-treated with increasing concentrations of Tg for 30 minutes prior to infection with HCoV-229E. The effect of Tg on viral replication and viral titer was assessed 48 hours later. Tg dose-dependently decreases both HCoV-229E intracellular N gene expression **(Figure 1B)** and viral titer **(Figure 1C).** Additionally, A549 cells were primed with Tg and infected with a HCoV-229E luciferase reporter virus to determine the effect of Tg on viral replication and cell viability **(Figure 1D)**. We show that Tg potently inhibits viral replication with a half-maximal inhibitory concentration (IC_50_) of 5 nM, as measured by luciferase reporter activity, without affecting cell viability at the tested concentrations **(Figure 1D).** To identify the replication step(s) that are inhibited by Tg, we performed a time of addition assay, in which A549 cells were treated with Tg for 30 min at various time points post-infection **(Figure 1E).** Tg most effectively inhibits viral N gene expression when added prior to 8-12 hours post-infection **(Figure 1E)**. Since transcription and RNA replication is expected to begin in this time window^14^, we investigated the effect of Tg on the accumulation of HCoV-229E transcripts and RNA replication intermediates. Tg treatment blocks viral gene expression **(Figure 1F),** completely inhibiting the accumulation of dsRNA in infected cells at 24 hours post infection **(Figure 1G).** These findings suggest that Tg inhibits HCoV-229E by blocking early stages of viral RNA replication at a post-entry step.

**Figure 1.**
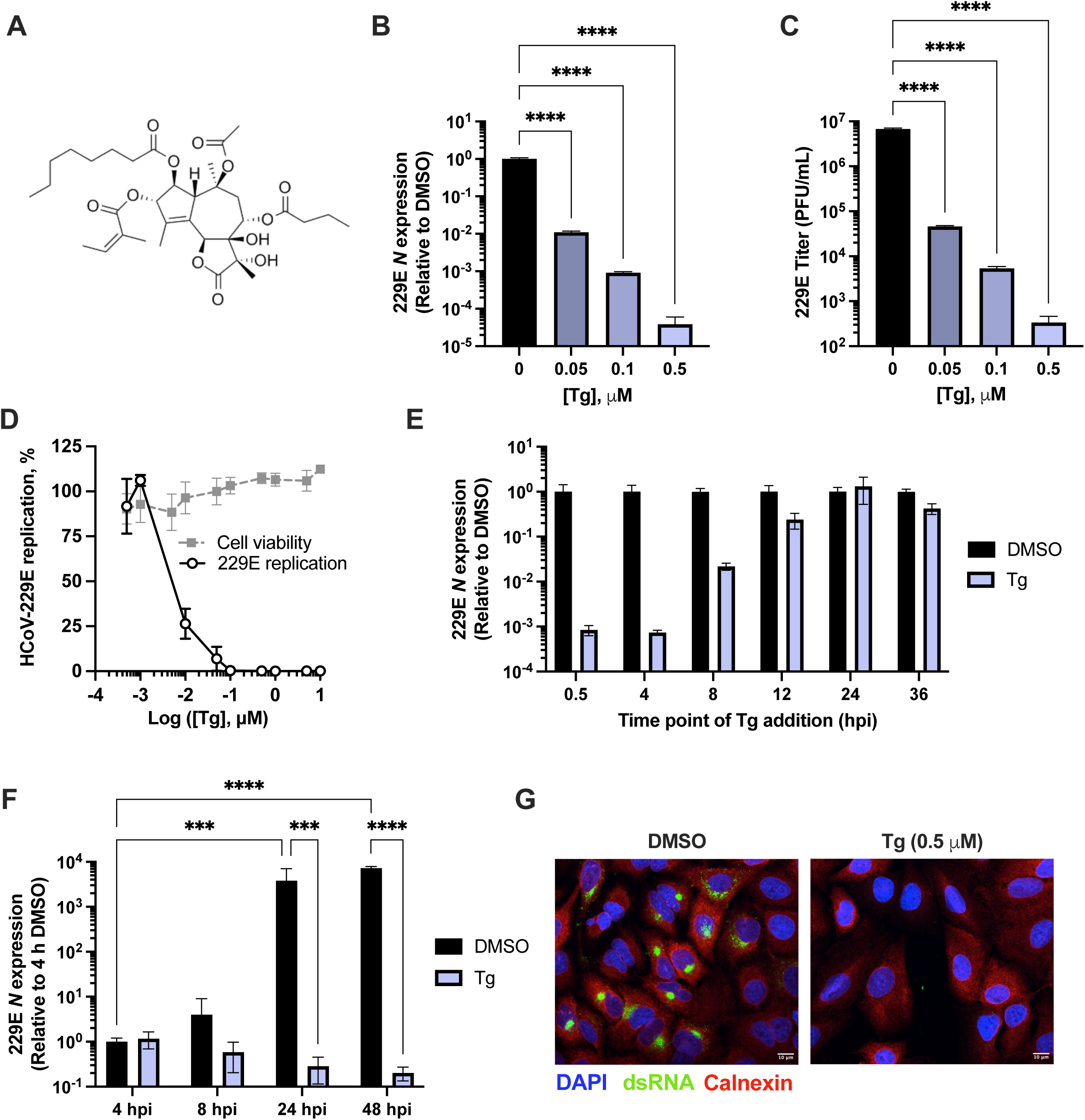
Tg inhibits HCoV-229E infection during early stages of RNA transcription/replication. **(A)** Chemical structure of thapsigargin. **(B-D)** A549 cells were primed with increasing concentrations of Tg for 30 minutes prior to infection with **(B-C)** HCoV-229E (MOI 0.01) or **(D)** a HCoV-229E luciferase reporter virus (MOI 0.5). **(B-C)** RNA lysates and viral supernatants were collected 48 hours post-infection (hpi). Viral supernatants were used to assess changes in viral titer via plaque assay. **(D)** Luciferase activity was read 48 hpi as a proxy for viral replication. Alamar blue assay was used to determine the cytotoxicity of Tg in A549 cells. **(E)** A549 cells were primed with Tg (0.5 μM) for 30 minutes at the indicated time points post infection. RNA lysates were collected at 48 hpi. **(F)** A549 cells were primed with Tg (0.5 μM) for 30 minutes prior to infection with HCoV-229E (MOI 0.05). RNA lysates were collected at the indicated time points post-infection. **(B, E, F)** RNA lysates were used to assess changes in gene expression by RT-qPCR. Data are normalized to *18S* and are expressed relative to DMSO. Graphs show means +/− SEM from three independent experiments performed in triplicate (***p<0.001, ****p<0.0001). **(G)** A549 cells were primed with DMSO or Tg (0.5 μM), then infected with HCoV-229E (MOI 0.5). At 24 hpi, cells were fixed and processed for immunofluorescence, staining for dsRNA (green) and the ER marker calnexin (red) using the appropriate antibodies.

### Thapsigargin expedites activation of the UPR during HCoV-229E infection

Tg potently induces ER stress and activates the UPR, which is comprised of three distinct signalling pathways, each initiated by a corresponding ER resident transmembrane protein (IRE1, PERK, or ATF6) **(Figure 2A)**. To confirm that Tg activates all three branches of the UPR in A549 cells, we primed cells with Tg for 30 minutes, and assessed expression of UPR-associated genes, representative of each pathway, over time **(Figure 2B)**. Induction of *XBP1*, *CHOP*, and *HERPUD1* gene expression were used as proxies for IRE1, PERK, and ATF6 pathway activation, respectively. Tg robustly activates each pathway at earlier time points, and although this activation wanes over time, expression of ER stress-associated genes was still elevated after 48 h in Tg-primed cells **(Figure 2B)**.

**Figure 2.**
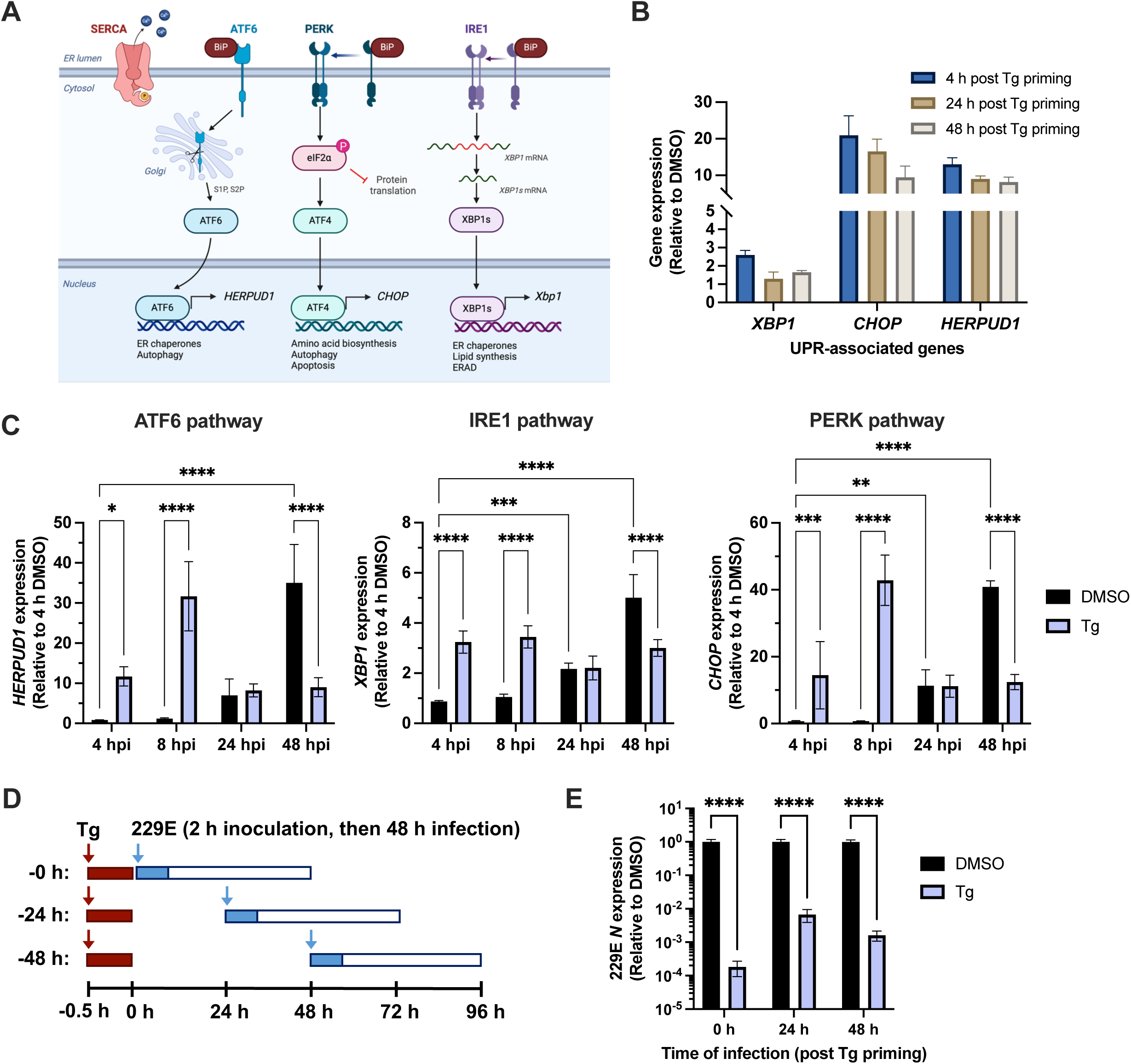
Thapsigargin broadly activates the UPR and expedites its activation during HCoV-229E infection. **(A)** Schematic of the ATF6, PERK, and IRE1 arms of the UPR. **(B)** Cells were primed with Tg (0.5 μM) for 30 minutes, and RNA lysates were collected at 4, 24 or 48 h after Tg priming. Expression of UPR-associated genes was assessed by RT-qPCR). **(C)** A549 cells were primed with DMSO or Tg (0.5 μM) for 30 minutes, then infected with HCoV-229E. RNA lysates were collected at 4, 24, or 48 hours after Tg priming. **(D-E)** Following 30 min of Tg priming, A549 cells were infected with HCoV-229E (MOI 0.05) immediately (0 h), or after 24 or 48 h. RNA lysates were collected at 48 h post-infection. **(B-E)** Changes in gene expression were assessed by RT-qPCR. Data are normalized to cellular *actin* and set relative to DMSO. Graphs show mean +/− SEM from three independent experiments performed in duplicate (*p<0.05, **p<0.01,***p<0.0005, ****p<0.0001).

During the course of infection, HCoV-229E infection induces ER stress **(Figure 2C)**, likely due to the manipulation of ER membranes to form viral ROs and the excessive translation and processing of viral glycoproteins. Although CoVs have evolved mechanisms to regulate ER stress responses^15^, we hypothesized that induction of ER stress by Tg disrupts the ability of HCoV-229E to regulate these responses. To characterize how Tg affects UPR activation in the context of infection, A549 cells were primed with Tg for 30 minutes, and subsequently infected with HCoV-229E. UPR-associated gene expression was assessed at various time points post-infection with or without Tg priming **(Figure 2C).** UPR-associated gene expression is induced earlier in infection (4 and 8 hpi) in cells primed with Tg in comparison to DMSO-treated cells, where infection induces the UPR at later time points and as late as 48 hpi **(Figure 2C).** *CHOP* expression, downstream of PERK pathway activation, was induced most strongly, with an approximate 40-fold increase in expression observed in A549 cells primed with Tg, relative to DMSO. These results suggest that Tg expedites the activation of the UPR during HCoV-229E infection, which may contribute to its antiviral effect.

Since we noticed a prolonged upregulation of UPR-associated genes following Tg priming, we next tested the duration of its antiviral effect. A549 cells were primed with Tg as described above, and subsequently infected with HCoV-229E at 0 h, 24 h or 48 h post-priming **(Figure 2D-E).** Remarkably, Tg retains its antiviral activity and significantly inhibits viral N gene expression, even when cells were infected 48 hours after Tg exposure **(Figure 2E)**, indicating that the inhibitory effect of Tg is long lasting.

### Inhibition of the IRE1 and ATF6 pathways does not impair the antiviral activity of Tg against HCoV-229E

We next tested whether any specific pathway(s) of the UPR mediate the antiviral effect of Tg against HCoV-229E infection, using pharmacological UPR inhibitors to selectively inhibit each pathway during human CoV infection. KIRA6^16,17^, GSK2606414^17^, and Ceapin-A7^18^ have previously been shown to selectively inhibit the IRE1, PERK, and ATF6 pathways, respectively. Unfortunately, the PERK inhibitor GSK2606414 was cytotoxic in A549 cells at all tested concentrations, so we focused on the IRE1 and ATF6 pathways. Treatment with KIRA6 reduced cell viability as measured by Alamar blue by approximately 50% in A549 cells, while Ceapin-A7 did not significantly affect cell viability, relative to DMSO (**Figure 3A)**. However, no overt morphological signs of cytotoxicity were observed by light microscopy, suggesting that KIRA6 may modulate cellular metabolic activity, or cell proliferation, rather than overtly inducing cell death. We first confirmed that KIRA6 and Ceapin-A7 inhibit the IRE1 and ATF6 pathway, respectively, by testing their ability to inhibit Tg-induced XBP1s and HERPUD1 protein expression (**Figure 3B-C)**. To determine the role of the IRE1 pathway in HCoV-229E infection and whether it mediates the antiviral effect of Tg, A549 cells were primed with DMSO or Tg alone, or in combination with KIRA6, and then infected with HCoV-229E and incubated in the presence of KIRA6. Co-treatment with KIRA6 significantly inhibits Tg-induced *XBP1s* expression during infection **(Figure 3D).** Interestingly, treatment with KIRA6 alone strongly inhibited HCoV-229E replication, making it difficult to assess any further antiviral effect of Tg **(Figure 3E-F)**. Nonetheless, these findings suggest that activation of the IRE1 pathway by Tg likely does not mediate its antiviral effect against HCoV-229E infection.

**Figure 3.**
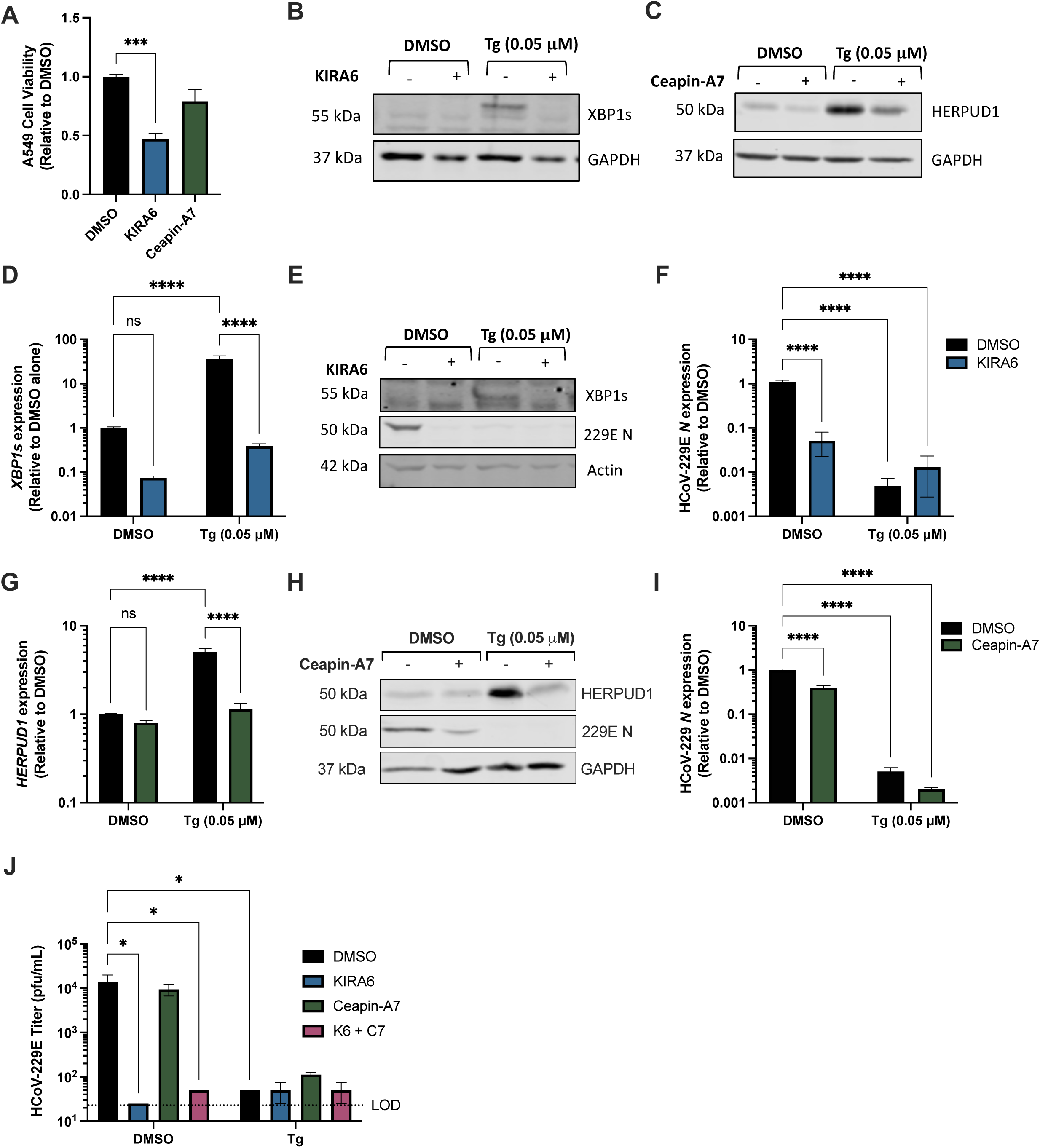
Pharmacological inhibition of IRE1 or ATF6 does not affect the antiviral activity of Tg against HCoV-229E infection. **(A)** A549 cells were treated with DMSO, KIRA6 (10 μM), or Ceapin-A7 (6 μM) for 24 hours. Alamar blue assay was used to assess cell viability based on cellular metabolic activity. **(B-C)** A549 cells were primed with DMSO or Tg (0.05 μM) in the absence or presence of KIRA6 (10 μM) or Ceapin-A7 (6 μM). Cell lysates were collected at 24 hours post-treatment. Xbp1s **(B)** or HERPUD1 **(C)** protein expression was assessed by western blot. **(D-F)** A549 cells were primed with DMSO or Tg (0.05 μM) in the presence or absence of KIRA6 (10 μM), prior to infection with HCoV-229E (MOI 0.05). Viral inoculum was removed 2 hours later and replaced with media containing DMSO or KIRA6 (10 μM). **(D-F)** Cell lysates were collected at 24 hpi to assess *XBP1s* and *N* gene expression by RT-qPCR **(D, F)**, or Xbp1s and HCoV-229E N protein expression by western blot **(E)**. **(G-I)** A549 cells were primed with DMSO or Tg (0.05 μM) in the presence or absence of Ceapin-A7 (6 μM) prior to infection with HCoV-229E (MOI 0.05). Viral inoculum was removed 2 hours later and replaced with media containing DMSO or Ceapin-A7 (6 μM). **(G-I)** Cell lysates were collected at 24 hpi to assess *HERPUD1* and *N* by RT-qPCR **(G, I)**, or HERPUD1 and HCoV-229E N protein expression by western blot **(H)**. RT-qPCR data are normalized to actin and set relative to DMSO. **(J)** A549 cells were primed with DMSO or Tg (0.05 μM) in the presence or absence of KIRA6 (10 μM) or Ceapin-A7 (6 μM), alone or in combination, prior to infection with HCoV-229E (MOI 0.05). Inoculum was removed 2 hours later and replaced with media containing the respective inhibitor(s). Viral supernatants were collected 24 hpi and titrated by plaque assay on Huh7 cells. Graphs show mean +/− SEM from three independent experiments performed in triplicate (*p<0.05, ****p<0.0001). Representative western blots are shown.

To assess the role of the ATF6 pathway, A549 cells were primed with DMSO or Tg in combination with Ceapin-A7 prior to infection **(Figure 3G-I)**. Ceapin-A7 inhibits Tg-induced HERPUD1 expression during infection, demonstrating its ability to inhibit ATF6 **(Figure 3G-H).** However, although Ceapin-A7 alone weakly inhibits HCoV-229E N expression, it does not impair the antiviral effect of Tg against HCoV-229E N gene or protein expression **(Figure 3H-I)**, ruling out a role for ATF6 pathway activation in the antiviral effect of Tg. Furthermore, co-treatment of infected cells with either KIRA6 or Ceapin-A7 alone, or in combination did not impair the ability of Tg to reduce viral titer **(Figure 3J)**, further confirming that the activation of these pathways does not contribute to the antiviral activity of Tg against HCoV-229E infection. Consistent with its effects on viral gene and protein expression **(Figure 3E-F)**, treatment with KIRA6 alone strongly reduced HCoV-229E titer **(Figure 3J)**.

### The antiviral activity of Tg against HCoV-229E is not dependent on IRE1, ATF6, or PERK expression

As a complementary approach to validate the role of each UPR pathway in mediating the antiviral activity of Tg, we generated stable knockdown A549 cells in which the expression of IRE1, PERK or ATF6 was silenced using RNAi **(Figure 4A)**. A549 shCTRL, shIRE1, shATF6, or shPERK cells were primed with DMSO or Tg and subsequently infected with HCoV-229E (MOI 0.05). The antiviral effect of Tg against HCoV-229E was not impaired in any of the knockdown cell lines **(Figure 4B-F)**, suggesting that Tg does not require the expression of any of these UPR sensors individually to inhibit HCoV-229E infection. Reflecting our earlier observations with the IRE1 inhibitor KIRA6 **(Figure 3E-F)**, we observed that HCoV-229E N protein expression was reduced in the shIRE1 cells in the absence of Tg treatment **(Figure 4B, Supp. Figure 1A)**, although there were no significant differences at the mRNA level **(Figure 4E)**. Tg priming enhanced IRE1 protein expression in the shIRE1 cells **(Figure 4B, Supp. Figure 1A)**, making it difficult to verify the importance of IRE1 in mediating the antiviral effect of Tg. Silencing of ATF6 reduces viral protein expression **(Figure 4C, Supp Figure 1B)**, reflecting our findings with Ceapin-A7 (the selective ATF6 pathway inhibitor), where Ceapin-A7 treatment reduces viral gene and protein expression **(Figure 3H-I).** In contrast to our results in the IRE1 and ATF6 knockdown cell lines, HCoV-229E N gene and protein expression is enhanced in the shPERK cells relative to shCTRL **(Figure 4D-E, Supp Figure 1C).** These data suggest that PERK activation inhibits HCoV-229E gene and protein expression. Surprisingly, viral titers were reduced by approximately 60% in each of the UPR knockdown cell lines.

**Figure 4.**
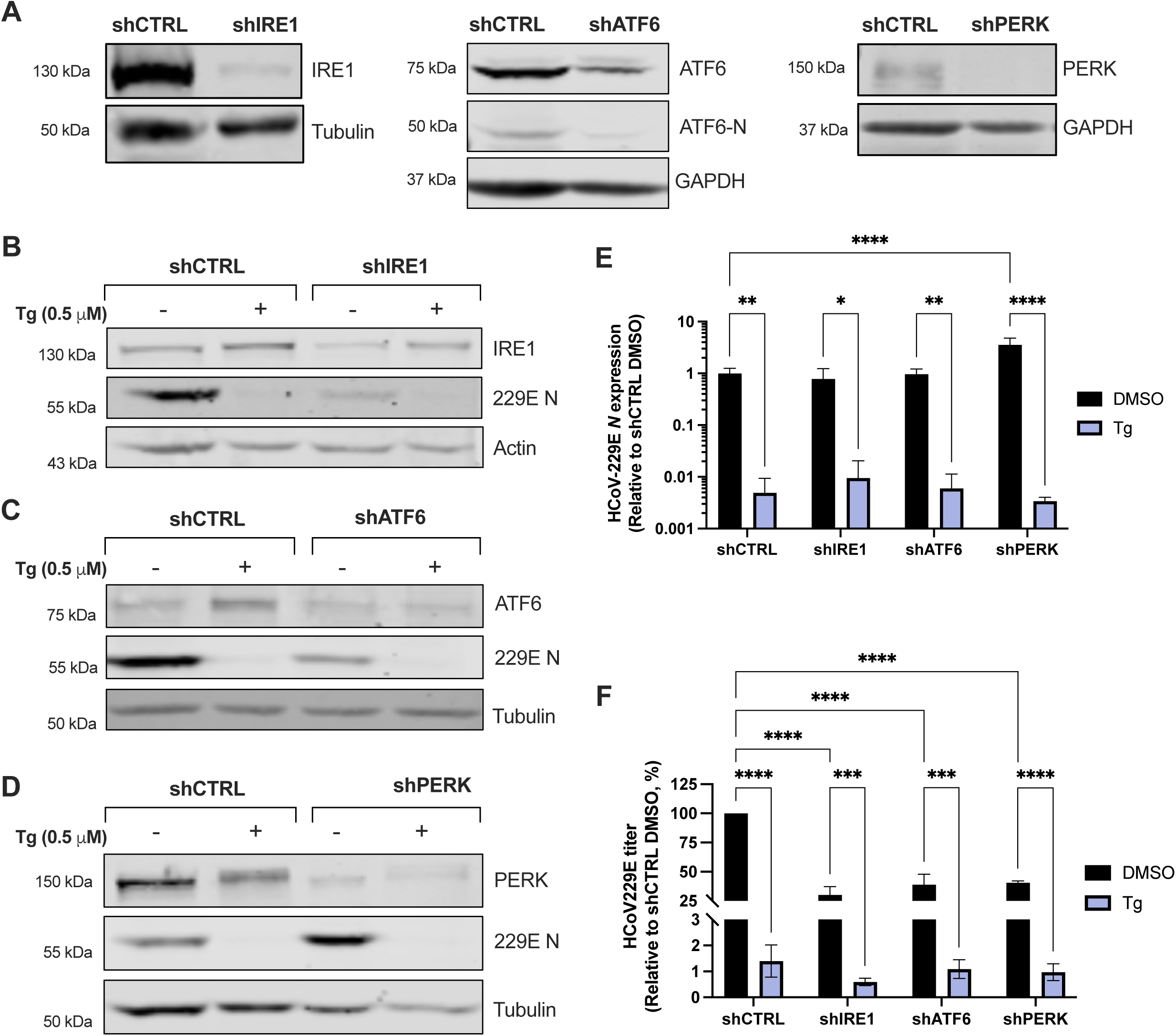
The antiviral effect of Tg against HCoV-229E infection does not require IRE1, ATF6, or PERK expression. **(A)** Stable UPR knockdown A549 cell lines were generated via lentiviral transduction. Silencing of protein expression was confirmed via western blot. **(B-F)** A549 shCTRL, shIRE1, shATF6, and shPERK cells were primed with DMSO or Tg for 30 minutes prior to infection with HCoV-229E (MOI 0.05). Cell lysates **(B-E)** or supernatants **(F)** were collected at 24 hpi. **(B-D)** Protein expression of IRE1, ATF6, PERK and HCoV-229E N were assessed by western blot. (E) Changes in gene expression were quantified by RT-qPCR. Data are normalized to actin and set relative to shCTRL DMSO. **(F)** Supernatants were used to determine viral titers by plaque assay on Huh7 cells. Due to variability between replicates, titration data is expressed as a percentage relative to the shCTRL DMSO in each experiment. Graphs show means +/− SEM from 3 independent experiments (*p<0.05, **p<0.01, ***p<0.001, ****p<0.0001).

### Selective activation of the PERK pathway inhibits HCoV-229E infection

As a complementary approach, we next used selective pharmacological UPR activators to determine whether activation of individual UPR pathways mimics the antiviral effect of Tg priming. IXA4^19^, AA147^20^, and CCT020312^21^ have previously been shown to selectively activate the IRE1, ATF6, and PERK pathways of the UPR, respectively. We confirmed activation of the respective UPR pathway by treating A549 cells with IXA4, AA147 or CCT020312 for 24 hours. *XBP1s* **(Figure 5A)**, *HERPUD1* **(Figure 5B)**, and *CHOP* **(Figure 5C)** mRNA expression was increased as expected in cells treated with the respective activators. Treatment of A549 cells with IXA4 or AA147 had no effect on cell viability relative to DMSO **(Figure 5D)**, although treatment of cells with CCT020312 or all three activators in combination somewhat reduced cell metabolic activity as assessed by Alamar blue assay **(Figure 5A)**. However, no overt signs of cytotoxicity were observed by light microscopy and cells were normal in appearance.

**Figure 5.**
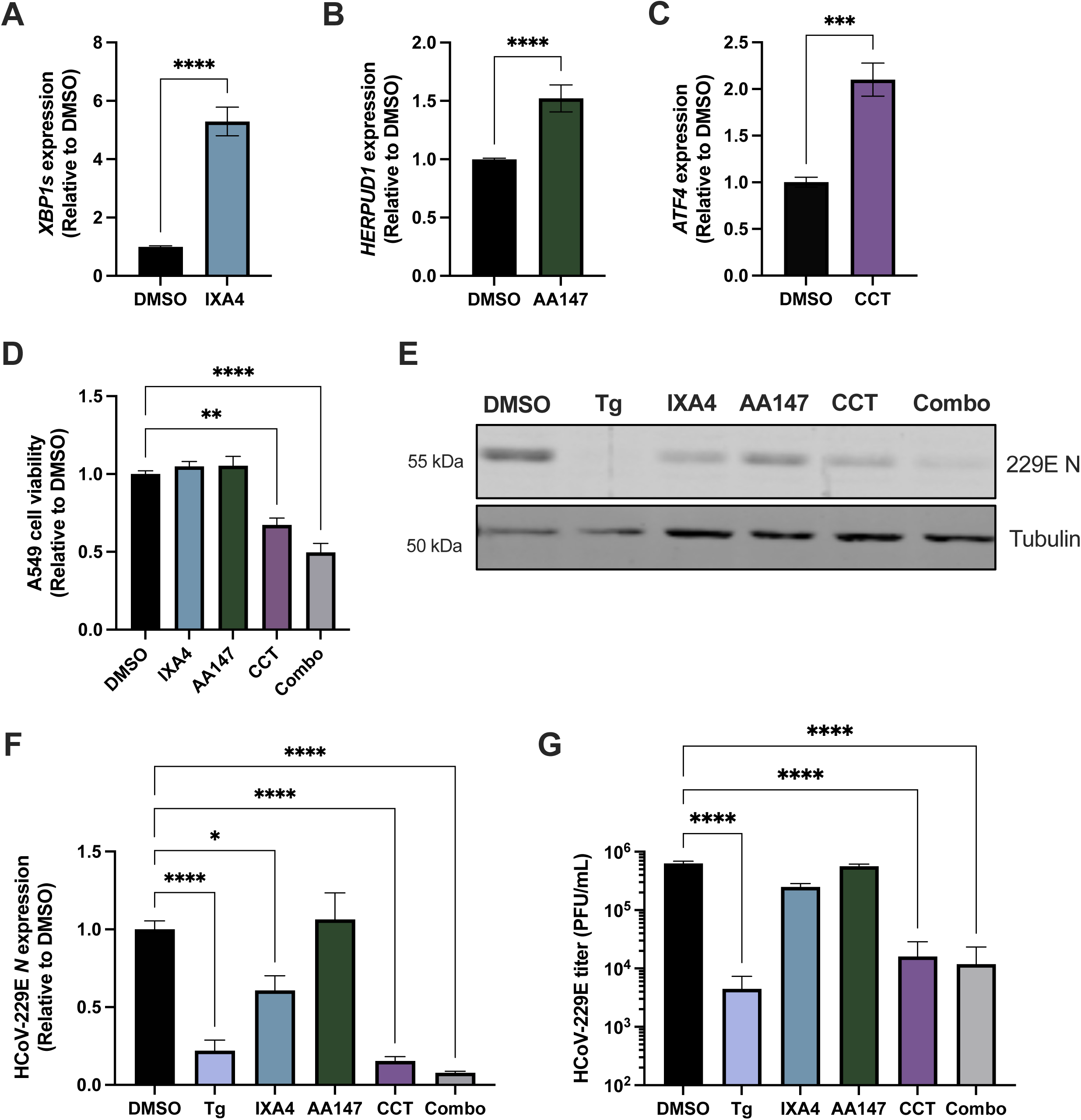
Selective pharmacological activation of PERK inhibits HCoV-229E replication. **(A-C)** A549 cells were treated with DMSO or one of the selective UPR activators for 24 hours. IXA4 (10 μM) selectively activates IRE1, AA147 (10 μM) selectively activates ATF6 and CCT020312 (5 μM) selectively activates PERK. RNA lysates were collected 24 hours post-treatment and the expression of genes downstream of each UPR pathway was assessed by RT-qPCR. Data are normalized to actin and set relative to DMSO. **(D)** Alamar blue assay was used to assess cell viability based on cellular metabolic activity in cells treated with the activators as described above. **(E-G)** A549 cells were primed with DMSO or Tg (0.05 μM) for 30 minutes prior to infection with HCoV-229E (MOI 0.05). Alternatively, cells were infected and then treated with media containing DMSO or the selective UPR activators alone or in combination. Cell lysates and supernatants were collected at 24 hpi. **(E)** Changes in HCoV-229E N protein expression were assessed by western blot. **(F)** Changes in HCoV-229E *N* gene expression was assessed by RT-qPCR. Data are normalized to actin and set relative to DMSO. **(F)** Viral titer was assessed by plaque assay. Data are relative to DMSO. Graphs show means +/− SEM or SD from 3 independent experiments (*p<0.05, ***p<0.01, ****p<0.0001).

To test effects of the UPR activators on HCoV-229E infection, A549 cells were primed with DMSO or Tg for 30 minutes or treated with the UPR activators for 24 hours following infection with HCoV-229E **(Figure 5E-G).** A549 cells treated with IXA4 show a mild reduction in HCoV-229E N gene and protein expression relative to DMSO, although this effect is minimal compared to Tg **(Figure 5E-F)**. IXA4 treatment does not significantly affect viral titer **(Figure 5G)**, further suggesting that activation of the IRE1 pathway likely does not mediate the potent antiviral effect of Tg. Although treatment of A549 cells with AA147 induces *HERPUD1* expression, it had minimal effect on HCoV-229E N gene expression **(Figure 5E-F)** or viral titer **(Figure 5G),** suggesting that activation of ATF6 does not underlie the antiviral activity of Tg. Interestingly, activation of PERK by CCT020312 significantly reduces HCoV-229E N gene and protein expression **(Figure 5E-F)**, as well as viral titer **(Figure 5F-G)** and recapitulates the antiviral effect of Tg. These results suggest that selective activation of PERK may at least partially underlie the antiviral effect of Tg. However, the simultaneous activation of all three pathways of the UPR, which occurs with Tg treatment, is likely required for robust inhibition of HCoV-229E infection.

### Selective activation of the PERK pathway of the UPR inhibits SARS-CoV-2 RNA replication

Since we showed that Tg priming inhibits CoV RNA replication **(Figure 1)**, we used a replicon model to evaluate whether selective activation of the PERK pathway could inhibit the replication of a more pathogenic coronavirus, SARS-CoV-2. We used BHK-21 cells stably expressing a SARS-CoV-2 replicon encoding a nanoluciferase reporter (BHK-CoV-2 cells)^12^. BHK-CoV-2 cells were primed with DMSO or Tg, or treated with CCT020312. RNA lysates were collected 24 hours later. Treatment with either Tg or CCT020312 significantly upregulated *ATF4* gene expression, confirming activation of the PERK pathway **(Figure 6A)**, without affecting cell viability **(Figure 6B)**. Cells treated with either Tg or CCT020312 both show a significant reduction in luciferase activity, a proxy for viral RNA replication, as well as SARS-CoV-2 *ORF1a* gene expression **(Figure 6C-D)**. Thus, selective activation of the PERK pathway of the UPR inhibits SARS-CoV-2 RNA replication, similarly to Tg. Furthermore, BHK-CoV-2 cells primed with DMSO or Tg, or treated with CCT020312, were processed for confocal microscopy and stained for double-stranded RNA (dsRNA), a viral replication intermediate. Consistently, BHK-CoV-2 cells treated with Tg or CCT020312 have decreased amounts of dsRNA **(Figure 6E)**. These findings show that selective activation of PERK significantly inhibits SARS-CoV-2 RNA replication, suggesting that PERK activation is broadly antiviral against human coronavirus infection.

**Figure 6.**
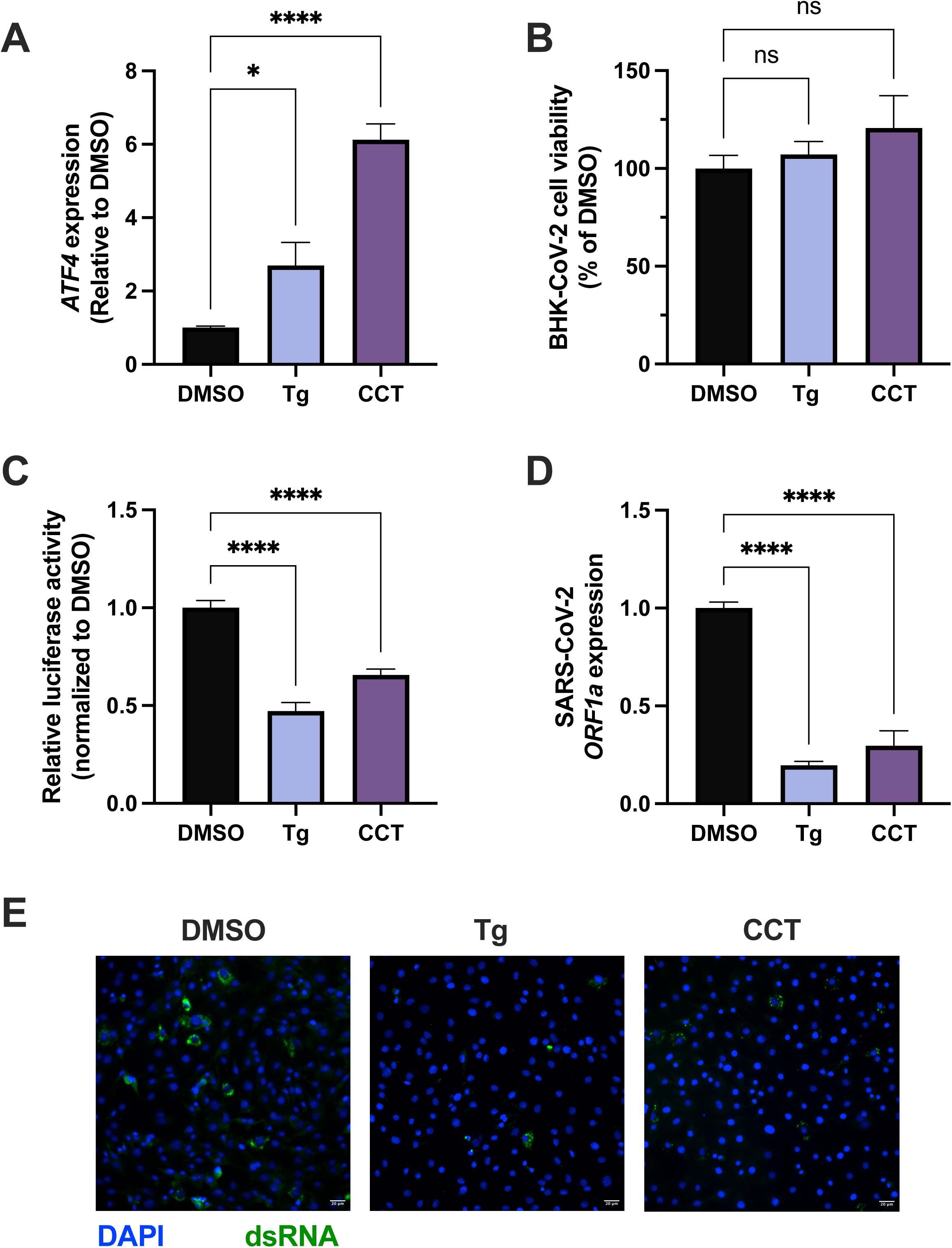
Selective activation of PERK inhibits SARS-CoV-2 RNA replication. **(A-E)** BHK-CoV-2 replicon cells were pre-treated with DMSO or Tg (0.5 μM) for 30 minutes, or treated with CCT020312 (5 μM) for 24 h, at which point *ATF4* mRNA expression **(A)**, cell viability **(B)**, and SARS-CoV-2 RNA replication **(C-D)** were assessed. **(A, D)** RNA lysates were collected 24 hours post-treatment. Data are normalized to hamster *GAPDH* and set relative to DMSO. **(E)** BHK-CoV-2 cells were treated with DMSO or Tg (0.5 μM) for 30 minutes, or CCT020312 (5 μM) for 24 hours. Cells were fixed 24 hours later and processed for fluorescence microscopy using an antibody for dsRNA (green). Cells were also stained for DAPI (blue). Graphs show means +/− SEM from 3 independent experiments (*p<0.05, ****p<0.0001).

## Discussion

Tg is a potent inducer of ER stress and the UPR, and possesses broadly acting antiviral activity against several unrelated viruses. While literature has shown that Tg inhibits both endemic (HCoV-229E, HCoV-OC43) and pathogenic (MERS-CoV, SARS-CoV-2) human coronaviruses^2,22^, its antiviral mechanism is not fully understood. In this study, we sought to characterize the antiviral mechanism of Tg and explore the role of the UPR in mediating the antiviral effect of Tg, using HCoV-229E as a model.

Our data demonstrate that a short 30-minute priming of A549 cells prior to infection with HCoV-229E robustly inhibits viral replication and abrogates the accumulation of viral dsRNA in infected cells at 24 hours post-infection. Furthermore, our time-of-addition experiments show that a 30-minute treatment with Tg significantly inhibits viral N gene expression when added within the first 8 hours of infection, suggesting that Tg exerts its antiviral effect against HCoV-229E infection by antagonizing early viral RNA synthesis. Like other +ssRNA viruses, CoV RNA transcription and replication occurs within cytoplasmic double-membrane replication organelles (ROs), the formation of which is mediated by viral non-structural proteins (Nsps)^23^. The timing of inhibition of viral replication by Tg is consistent with the timing at which the translation of key viral proteins involved in the formation of the CoV replication-transcription complex take place, and the abrogation of dsRNA accumulation in Tg primed cells suggests that the early stages of RNA synthesis are impaired.

Since CoV infection itself can induce ER stress and the UPR, we explored how Tg modulates the activation of each arm of the UPR during infection. Infection of A549 cells with HCoV-229E in the absence of Tg broadly activates all three pathways of the UPR, but this activation only occurs much later in infection (48 hours post infection). In contrast, expression of UPR-related genes in Tg-primed cells was significantly higher at earlier time points, indicating that Tg priming expedites the activation of the UPR in infected cells. Of all three representative UPR-associated genes tested, the mRNA expression of CHOP, downstream of PERK pathway activation, appeared to be most strongly upregulated. This strong induction of the PERK pathway is consistent with previous literature, showing that other coronaviruses, including SARS-CoV^24^ and MERS-CoV^25^, activate PERK and induce ATF4/CHOP expression. Activation of PERK by these viruses may serve a proviral role, as CoV infection can manipulate autophagy and ERAD machinery downstream of PERK pathway activation to support viral RO formation^26^. Additionally, betacoronaviruses, including SARS-CoV-2 and murine hepatitis virus (MHV) can induce autophagy to promote viral egress through the lysosomal pathway^27^.

To further explore the role of each UPR pathway in HCoV-229E infection and the effect of Tg, we used selective pharmacological UPR activators or inhibitors. KIRA6 and Ceapin-A7 were used to selectively inhibit the IRE1 and ATF6 arms of the UPR, respectively. Blockade of these pathways did not affect the ability of Tg to inhibit viral N gene or protein expression, suggesting that the activation of either of these pathways likely does not contribute to its antiviral effect against HCoV-229E. Consistently, selective activation of the IRE1 or ATF6 pathways did not significantly inhibit HCoV-229E infection, as measured by viral titer.

In contrast, selective activation of the PERK pathway using CCT020312 significantly inhibits HCoV-229E replication and most closely mimics the antiviral effect of Tg treatment. The antiviral effect of PERK pathway activation has been shown in the context of another alphacoronavirus, transmissible gastroenteritis virus, where PERK activation during infection negatively regulates viral replication^28^. Additionally, reflecting our observations in A549 cells infected with HCoV-229E in the absence of Tg, previous literature shows that infection with infectious bronchitis virus (an avian coronavirus) induces the expression of downstream effectors of PERK pathway activation, including CHOP, at 24-48 hpi^29^. Therefore, it is possible that skewing the kinetics of UPR and particularly PERK pathway activation to occur earlier during infection may antagonize viral RNA replication.

To validate the role of PERK pathway activation in mediating the antiviral effect of Tg against human coronavirus infection more generally, we assessed the ability of CCT020312 to inhibit replication of SARS-CoV-2 replicon RNA. This allowed us to explore the antiviral effect of CCT020312 against a more pathogenic and clinically relevant human CoV, while also providing the opportunity to test the antiviral effect of Tg and CCT020312 specifically against CoV RNA replication. As we observed in the context of HCoV-229E infection, treatment of BHK21-CoV-2 replicon cells with CCT020312 significantly inhibited SARS-CoV-2 RNA replication.

The PERK pathway plays a critical role in regulating host cellular translation through the phosphorylation of eIF2α, leading to the global attenuation of cap-dependent translation. Given that CoVs rely on host cap-dependent translation machinery to readily translate their large polyproteins upon entry into the cell, many CoVs have evolved strategies to evade host translation shutoff. For example, SARS-CoV-2 non-structural protein 1 (nsp1) can block the translation of host mRNA through interactions with the 40s ribosomal subunit. Viral mRNA can evade this inhibition and selectively mediate their translation via interactions between the 5’ untranslated region and host ribosomes^30,31^. However, it is still possible that activation of PERK by Tg during early infection suppresses the translation of viral polyproteins encoding key viral nsps. As these are required for the assembly of the replication-transcription complex, this could explain why priming of A549 cells with Tg significantly inhibits the transcription of viral mRNAs and the accumulation of viral dsRNA in infected cells. PERK activation is also associated with the upregulation of pro-apoptotic factors such as CHOP. In addition to protein translation halt, the activation of apoptosis by PERK may promote death of infected cells and limit viral spread. Alternatively, as mentioned above, PERK activation can trigger autophagy and selective ER-phagy^26^. This may impair coronavirus replication by degrading ER-derived membranes used for viral replication complex formation, or by targeting viral replication intermediates, such as dsRNA, for degradation.

While activation of the IRE1 pathway did not appear to contribute to the antiviral activity of Tg, we found that treatment of A549 cells with the selective IRE1 inhibitor KIRA6 strongly inhibits HCoV-229E infection. Consistently, previous literature shows that inhibition of IRE1 during SARS-CoV-2 infection inhibits S protein expression and significantly reduces viral titers in infected A549 cells^32^. Another paper showed that the inhibition of IRE1 using another selective inhibitor, KIRA8, decreases HCoV-OC43 N protein expression in A549 cells, although the replication efficiency of HCoV-OC43 and other beta coronaviruses tested did not appear to be impaired by KIRA8^33^. One functional consequence of IRE1/XBP1s activation is the induction of ER-associated degradation (ERAD) and the transcription of ERAD-associated genes. ERAD is a quality-control system that facilitates the retro-translocation of misfolded/unfolded proteins out of the ER into the cytosol, tagging them for proteasomal degradation^34^. Several CoVs upregulate ERAD components during infection^15,26^. Interestingly, MHV, a murine betacoronavirus, hijacks EDEMosomes, which are vesicles involved in the transport of ERAD machinery and client proteins from the ER to the lysosome, for use as ROs^26^. While additional experimentation is required to understand the role(s) of the IRE1 pathway during HCoV-229E infection, it is possible that human CoVs may similarly utilize ERAD machinery to facilitate their replication, which could explain why inhibition of IRE1/XBP1s activity reduces HCoV-229E infection. However, one previous paper showed that the activation of the IRE1 pathway and downstream XBP1 splicing was enhanced in Vero E6 cells infected with a SARS-CoV virus lacking the envelope (E) gene, suggesting that the E protein may play a role in antagonizing the IRE1 pathway^35^. These findings highlight the duality of the UPR in CoV infection, and reflect that these processes are tightly regulated during viral infection. Previous research investigating the role of the IRE1 pathway during coronavirus infection has predominantly focused on betacoronaviruses (SARS-CoV-2, HCoV-OC43, MHV). Our data identifies a similar role for the IRE1 pathway in supporting the replication of alphacoronaviruses.

Together, our data furthers the understanding of the antiviral mechanism of Tg against human coronavirus infection. We speculate that Tg-mediated activation of the PERK pathway drives its antiviral effect, since activation of PERK impairs HCoV-229E replication and reduces viral titer, and silencing of this arm of the UPR promotes viral replication. Activation of PERK during infection may impair the translation of viral RNA, thereby preventing the synthesis of key viral polyproteins and the formation of the replication-transcription complex. However, the precise mechanisms remain to be established. While silencing of PERK expression by shRNA did not impair the antiviral activity of Tg against HCoV-229E replication, it is possible that the combined activation of all three pathways by Tg contributes to its antiviral effect. Alternatively, residual expression of PERK in the shRNA cell line may be sufficient to mediate the antiviral effect of Tg. Future experiments in knockout cell lines, or with PERK inhibitors, would help to address this. Another possibility is Tg exerts its antiviral effect in a UPR-independent manner. Tg potently inhibits SERCA2, thereby disrupting calcium flux and broadly inducing ER stress, but it is possible that SERCA2 itself could act as a host factor for HCoV-229E, such that inhibition of SERCA2 by Tg inhibits viral infection, independently of the UPR. Previous studies have also implicated Tg in priming antiviral IFN responses against IAV^4^, as well as flaviviruses including dengue virus and Zika virus^36^. Thus, the antiviral effect of Tg against human CoV infection could be associated with heightened antiviral responses following Tg priming. Further experimentation is required to validate these hypotheses. Additionally, the use of A549 cells within our study, an adenocarcinoma-derived cell line, is a possible limitation as these cells may respond differently to ER stress. Therefore, in future studies it is important to validate these findings in primary human airway epithelial cells.

## Conclusion

In summary, our findings provide insight into the antiviral mechanism of Tg against human CoV infection. We show that Tg priming of A549 cells accelerates the activation of the UPR during HCoV-229E infection and inhibits early stages of viral RNA synthesis. However, the antiviral effect of Tg does not appear to be dependent on any individual UPR pathway. We show that activation of PERK, but not IRE1 or ATF6, inhibits HCoV-229E infection, with PERK activation also inhibiting SARS-CoV-2 RNA replication. Inhibition of IRE1 potently inhibits HCoV-229E infection, suggesting a proviral role for this arm of the UPR against human coronavirus infection. Overall, we have characterized the role of the UPR in HCoV-229E infection and furthered our understanding of the antiviral mechanism of Tg. These studies could inform the development of host-targeted antiviral strategies to prepare for future emerging CoVs.

## Data availability statement

All the data supporting this article have been included in the main text or in the supplementary information.

## Acknowledgements

This work was supported by the Natural Sciences and Engineering Research Council of Canada (CCC) and the Canadian Institutes for Health Research (CCC). We are grateful to the Canadian Foundation for Innovation John R. Evans Leaders Fund (CCC) for equipment that supported this project. THT acknowledges support from a Canada Graduate Scholarship – Master’s (CGS-M) and a Vanier Canada Graduate Scholarship. THT and CEGF acknowledge PhD studentships and support from the Canadian Network on Hepatitis C. We are grateful to Dr. Craig McCormick (Dalhousie University, Canada) for providing reagents and helpful discussion. We are grateful to Dr. Volker Thiel (University of Bern, Switzerland) for providing the HCoV-229E-RLuc reporter virus. The following reagent was obtained through BEI Resources, NIAID, NIH: BHK-21 Cell Line Harboring SARS-CoV-2-Replicon Containing NanoLuc^®^-Neo Reporters and NSP1 Mutations (K164A/H165A), NR-58876.

## Supplementary Figure Captions

**Supp. Figure 1. Silencing of UPR sensors and the effect on HCoV-229E protein expression**. **(A-C)** Band density analysis was performed in Fiji ImageJ and are expressed relative to the band density ration obtained in shCTRL DMSO condition. Representative western blots are shown in Figure 4B-D.

