## Supplementary figures and images for "Chemical modulation of the unfolded protein response reveals an antiviral role for the PERK pathway in human coronavirus 229E infection"

### Supplemental Figure 1

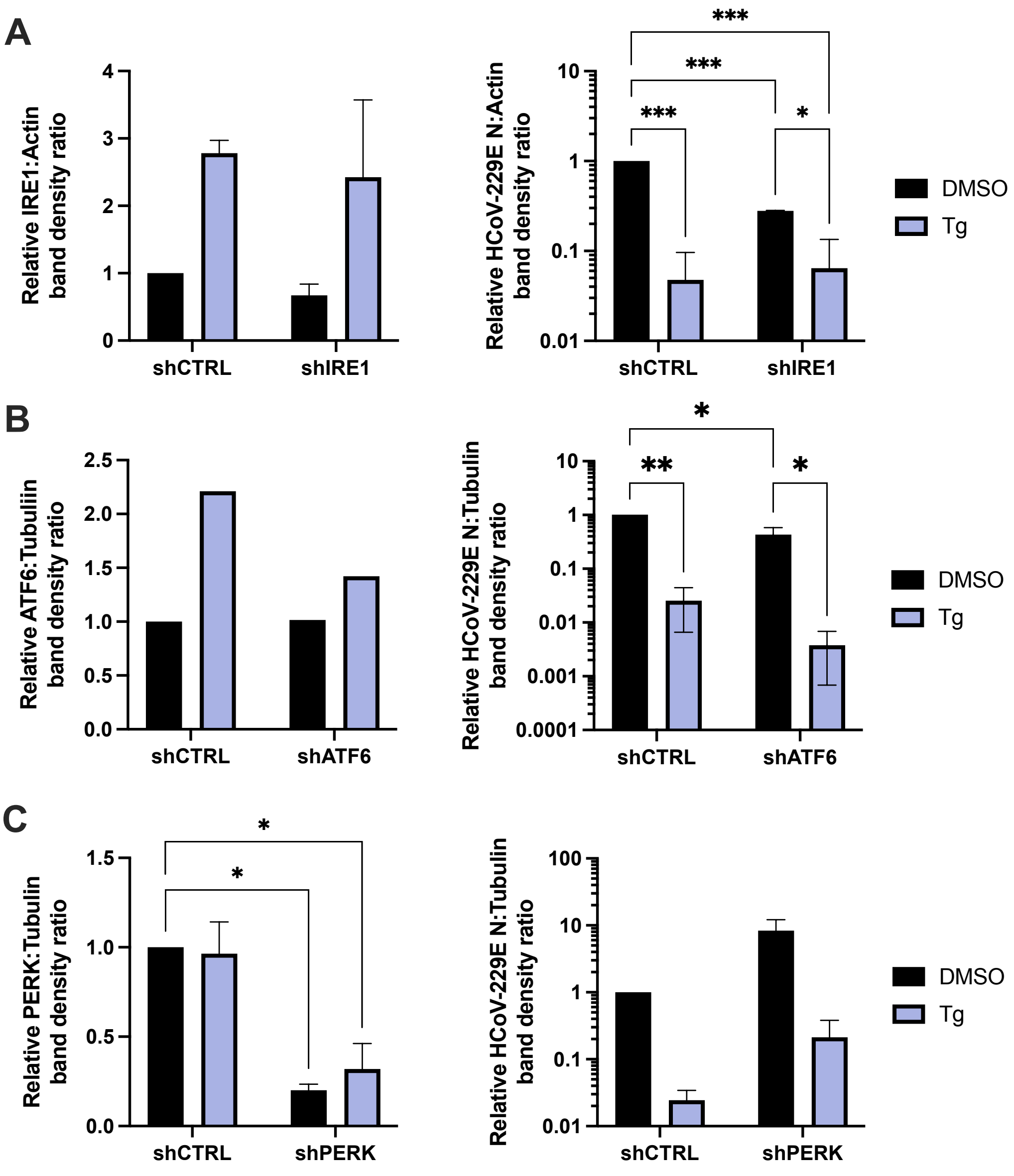
